# Seroprevalence of Human T-Lymphotropic Virus Type-1 (HTLV-1) Among Blood Donors at a Tertiary Hospital in Abakaliki, Ebonyi State, Nigeria

**DOI:** 10.64898/2026.06.09.731045

**Authors:** E. N. Onu, D. C. Ilang, C. O. Akpa, E. M. Onu, C. E. Orji, R. O. Ologwu, E. A. Obaji, L. O Nomeh

## Abstract

The study was a cross-sectional study done on the Blood Bank Unit of the Federal Teaching Hospital, Ebonyi State, Nigeria among blood donors.

**Methods:** The total number of prospective blood donors that were initially screened against HIV, Hepatitis B and Hepatitis C virus was 300, to which, 93 eligible donors were incorporated in the study. Anti-HTLV-1 IgM and IgG antibodies in serum were also tested in the Enzyme-Linked Immunosorbent Assay (ELISA) technique. The descriptive statistics were used in the analysis of data.

**Results:** Out of the 93 blood donors analyzed, 23 (24.6%) were positive for HTLV-1 antibodies. Seropositivity was highest among donors aged 30–35 years, with anti-HTLV-1 IgM and IgG prevalences of 35.3% and 29.4% respectively, compared with lower values in other age groups. Married donors showed higher IgM (32.8%) and IgG (34.1%) seroprevalence than single donors (IgM: 20.3%; IgG: 18.9%). Female donors recorded higher seroprevalence (IgM: 26.0%; IgG: 27.3%) than male donors (IgM: 23.4%; IgG: 22.1%). Occupational distribution showed the highest seroprevalence among artisans (IgM: 37.5%; IgG: 37.5%), followed by students (IgM: 25.7%; IgG: 22.9%), traders (IgM/IgG: 23.8%), and civil servants (IgM: 14.3%; IgG: 19.0%). Variance analysis revealed significant differences among serological patterns (IgG^−^IgM^+^, IgG^+^IgM^−^, IgG^+^IgM^+^, IgG^−^IgM^−^), with mean values differing significantly at p ≤ 0.05.

**Conclusion:** This research has revealed that seroprevalence of HTLV-1 was very high among the blood donors, which indicates that there is the possibility of transfusion-transmitted infection. Routine HTLV-1 screening as the measure to enhance blood safety in the area is important.

## Introduction

Human T-Lymphotropic Virus type-1 (HTLV-1) is a human retrovirus belonging to the family of Retroviridae and genus to Deltaretrovirus. It was highly visibly the original human retrovirus and is pathologically linked to a variety of debilitating disorders, including adult T-cell leukemia/ lymphoma (ATLL) and HTLV-1/myelopathy/tropical spastic paraparesis (HAM/TSP) [1,2]. Although most individuals who contract the infection do not manifest the infection in the course of their lifetime, HTLV-1 infection results in the viral persistence in people throughout their lifetime and is a severe concern of the community health in the endemic regions. HIV-1 is predominantly transmitted by means of transfusion of infected cell blood constituents, sex, mother to child particularly when prolonged breastfeeding as well as sharing needles and sharp objects [3,4]. Blood transfusion is quite prolific in the number of such ways particularly where unscreened blood is employed. It has been established that transfusion of cellular blood elements infected by HTLV-1 can cause seroconversion in up to 20-60 percent of its recipients [5,6].

It is approximated that there are about 5 to 10 million infected with HTLV-1 individuals in the world with endemic regions reported in the sections of Japan, Caribbean, South America, sub-Saharan Africa and Melanesia [7,8]. The extent of spread of HTLV-1 infection in Africa has been considerably extensive but the burden of the same has been underestimated due to the absence of surveillance activities as well as absence of routine screening of blood transfusion services in most countries in Africa. Blood donors in Nigeria are regularly screened with regards to Human Immunodeficiency Virus (HIV), Hepatitis B Virus (HBV), and Hepatitis C Virus (HCV) but in screening HIV, HBV and HCV, HTLV-1 has not been included in the Nigerian policy of blood transfusion policy. Nigeria has reported many studies with various prevalence of the HTLV-1 in blood donors and other population groups having low to moderate levels depending on the geography of the location where the study was carried out and the study design used [9,10,11]. Such findings point out to the continuous spread of HTLV-1 in the population and the possibility that there may be transfusion-transmitted infection. However, lack of information about the occurrence of HTLV-1 in blood donors in southeastern region of Nigeria is present.

The town of Abakaliki is the capital of the Ebonyi State that provides high population with a tertiary healthcare organisation called as Federal Teaching Hospital Abakaliki (FETHA) and is also the foremost referral centre in the region. Blood donation in these regions is highly critical since it involves patients, and therefore health safety of donated blood poses a health issue to citizens. This study was seeking to determine the seroprevalence and the serology distribution of the HTLV-1 infection amongst blood donors in a tertiary hospital in Abakaliki at Ebonyi State in Nigeria as there is no information on the region area and since screening of the HTLV-1 infection is not a routine practice in the country. The findings of this study will be successful in strengthening the existing body of information and can be applied in making policies related to blood safety and blood transfusion in Nigeria.

## Materials and Methods

### Study Centre

This research was carried out in the Blood Bank Unit of the Federal Teaching Hospital at Abakaliki (FETHA), Abakaliki, Ebonyi State, southeast of Nigeria. The hospital is a large tertiary healthcare hospital that is a referral centre to the Ebonyi State and other neighbouring states. In the south eastern area of Nigeria, Abakaliki is located in the tropical rain forest belt and the climate has not been an exception to the area since it is warm throughout the year.

### Ethical approval and informed consent

Ethical clearance with reference number (**AE-FUTHA/REC/VOL 7/ 2025/722)** was obtained from Alex-Ekwueme Federal University Teaching Hospital, Abakaliki. All participants were duly informed of the objectives of the study and the protocol for sample collection. Participation was voluntary

### Study population

Prospective blood donors who reported to the Blood Bank Unit of the Federal Teaching Hospital Abakaliki (FETHA), Abakaliki, Ebonyi State were used to obtain blood samples to test them. The blood bank gets replacement and voluntary donors on a daily basis. Consecutive recruitment of all willing blood donors was done over the study area. The Research and Ethics Committee of the Federal Teaching Hospital Abakaliki gave the study an ethical approval. The participants were helped to do the structured questionnaires aimed to receive the adequate information such as demographic features and the history of the prior blood donations and transfusion. The exclusion criteria were a history of chronic disease (i.e. hypertension or other chronic ailments) and age younger than 18 years or older than 59 years (the recommend age range to donate blood). Three hundred (300) potential blood donors were screened, recruited and positive to HIV, HBV and HCV before being included into the HTLV-1 study. The HIV-1/2tm test kit (Abbott Japan Co., Ltd., Tokyo, Japan) was used to screen HIV-1/2 and screening the Hepatitis B surface antigen (HBsAg) and the Hepatitis C virus antibodies was done using the rapid test kits (Abon Bio pharma, Hangzhou, China). Any donor who revealed positive results on any one of these tests was not allowed to continue his participation in the study.

### Collection of Samples and Analysis

Three hundred (300) blood donors were recruited and screened out of which ninety-three (93) of the donors qualified according to the study eligibility criteria. Four point five milliliters (5 ml) of venous blood was aseptically transferred to sterile plain sample bottles of each eligible blood donor. The blood samples were left to clot and later centrifuged at 3000 revolutions per minute (rpm) in 10 minutes. The individualised serums were transferred cautiously in sterile cryovials and kept at 20 degree centigrade until laboratory examination.

Human T-Lymphotropic Virus type-1 (HTLV-1) IgM and IgG serum samples were tested in the Enzyme-Linked Immunosorbent Assay (ELISA) method to test the presence of materials stored in serums. The antibodies that were specific to HTLV-1 were detected by means of a commercially available HTLV-1 IgM/IgG ELISA kit produced by WKEA Med Supplies Corporation (China) in compliance with the manufacturer protocols strictly. Enzyme immunoassays were carried out on all the specimens and the presence or absence of the antibodies of HTLV-1 IgM and IgG was tested by comparing the optical density (absorbance) values of the test samples with the absorbance value of the cut-off calibrator of the kit. The edited data following the laboratory examination were keyed in and run through the descriptive statistical analysis in R-studio

## Results

Figure 1 showed that among the 93 blood donors, the sera of which were examined by the methods of enzyme immunoassay (ELI), by the help of the complement-fixing IgM and IgG antibodies against the HTLV-1 virus, 24.6% or 23 of them were positive in both complement-fixing antibodies. As indicated in Table 1, the greatest prevalence of anti-HTLV-1-IgM was observed in the donors with the highest prevalence of 6 (35.3%) donors recording positive results whereas the highest prevalence of anti-HTLV-1-IgG was on the same group of donors where 5 (29.4%) donors gave positive results. This age group hence showed the highest level of exposure to HTLV-1 of the study population. The percentage of positive anti-HTLV-1 immunoglobulin M (IgM) and anti-HTLV-1 immunoglobulin G (IgG), was identified to be 4 (26.7) and 5 (33.3) in the group of donors aged 36-41 years, respectively. Those donors that fell in the age bracket (42-49 years) achieved 4 (30.8) positivity of anti-HTLV-1 IgM and IgG antibodies. In the younger age groups, the age group of 18-23 years were positively persistently (3 (15.0) and 4 (20.0)) and age group of 24-29 years were positively persistently (4 (21.1) and 3 (15.8)) in relation to anti-HTLV-1 IgM and anti-HTLV-1 IgG respectively. Two (22.2) out of the donors in the age-group of 50-59 years were positive on anti-HTLV-1 IgM; 2 (22.2) on anti-HTLV-1 IgG.

**Figure 1:**
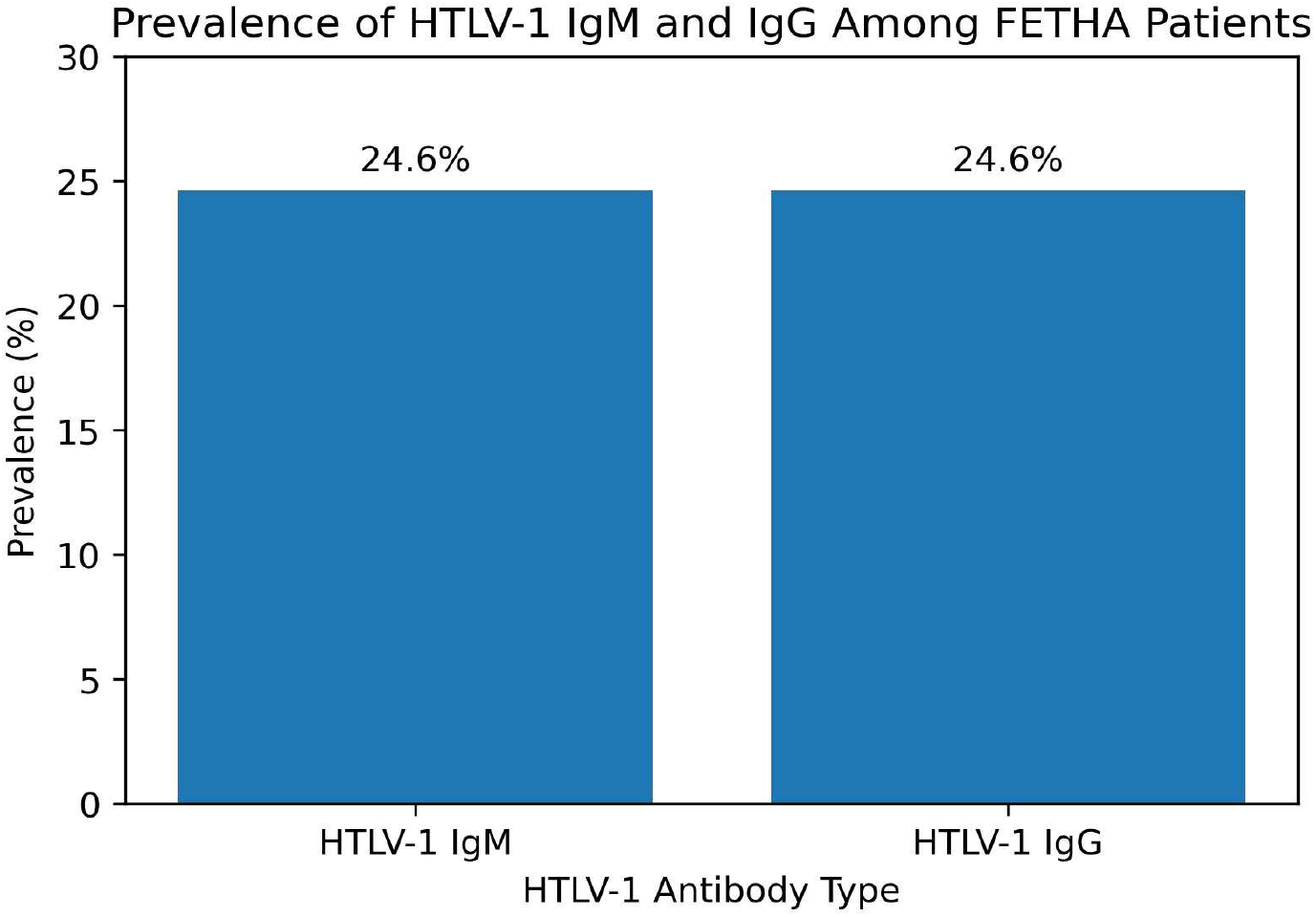
Prevalence of HTLV-1 IgM and lgG among FETHA blood donors.

**Table 1:** Age Distribution of HTLV-1 Antibodies Among the Study Subjects.

| Age Range<br>(years) | No. of Subjects<br>Tested (%) | No. of Subjects with anti-HTLV-1 IgM n (%) | No. of Subjects with anti-HTLV-1 IgG n (%) |
| --- | --- | --- | --- |
| 18 – 23 | 20 (21.5) | 3 (15.0) | 4 (20.0) |
| 24 – 29 | 19 (20.4) | 4 (21.1) | 3 (15.8) |
| 30 – 35 | 17 (18.3) | 6 (35.3) | 5 (29.4) |
| 36 – 41 | 15 (16.1) | 4 (26.7) | 5 (33.3) |
| 42 – 49 | 13 (14.0) | 4 (30.8) | 4 (30.8) |
| 50 – 59 | 9 (9.7) | 2 (22.2) | 2 (22.2) |
| **Total (%)** | **93 (100)** | **23 (24.6)** | **23 (24.6)** |

The figure 2 showed high prevalence of antibodies to HTLV-1 amongst the married blood donors compared to the single blood donors as indicated in the figure. In particular, the anti-HTLV-1 IgM was higher among married individuals (32.8) than single individuals (20.3). In the same case, the anti-HTLV-1 IgG antibodies were more prevalent among the married (34.1) as compared to the single (18.9) subjects.

**Figure 2:**
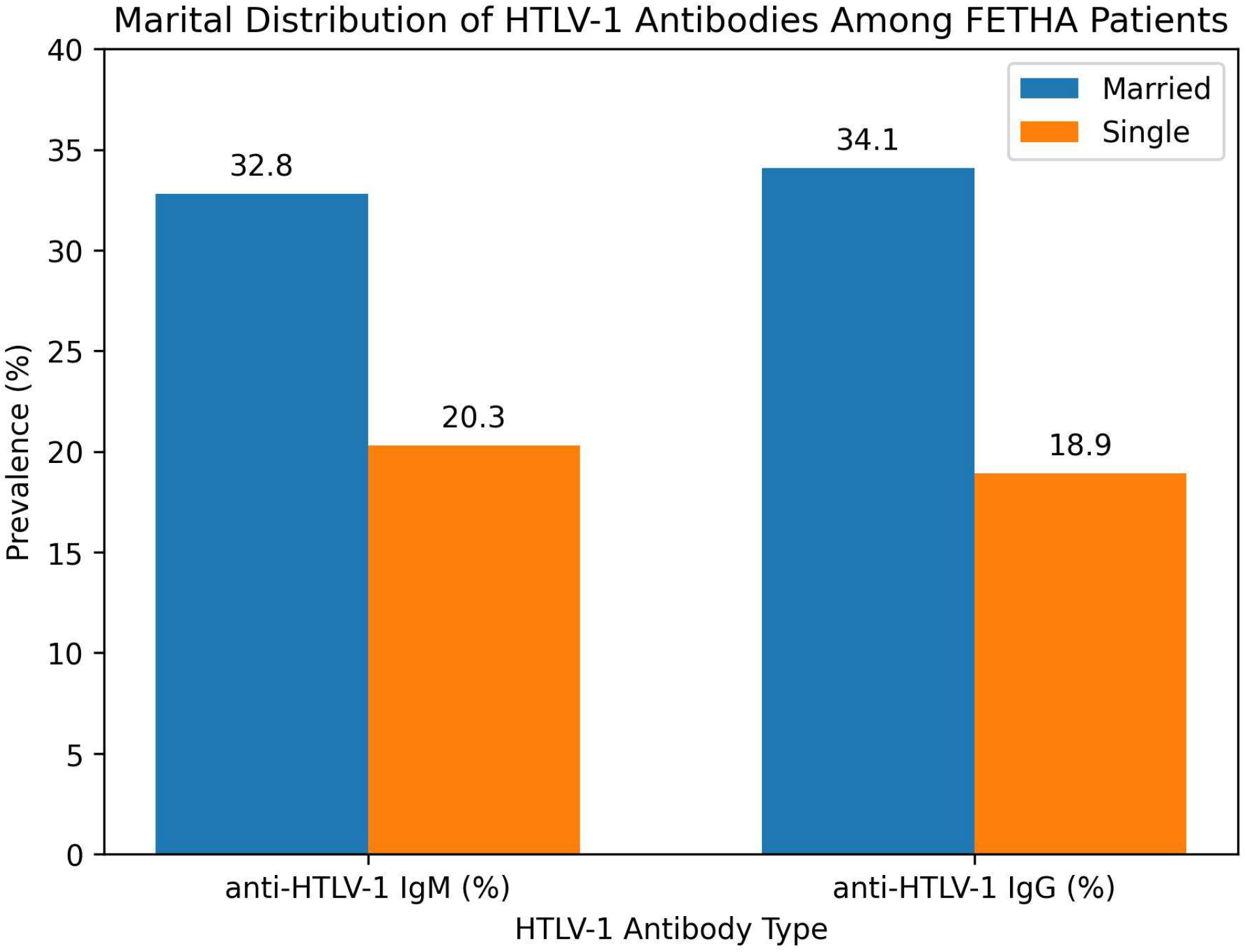
The illustration of the proportion of marital status on human T-Lymphotropic Virus type-1 (HTLV-1) antibodies among the study subjects

Figure 3 showed the sex proportion of the Human T-Lymphotropic Virus type-1 (HTLV-1) antibodies of the subjects in this study. The female subjects (26.0%) had a higher prevalence level of anti-HTLV-1 IgM compared to the male subjects (23.4%). On the same note, females were more prone to anti-HTLV-1 IgG antibodies as compared to males (27.3% vs. 22.1%).

**Figure 3:**
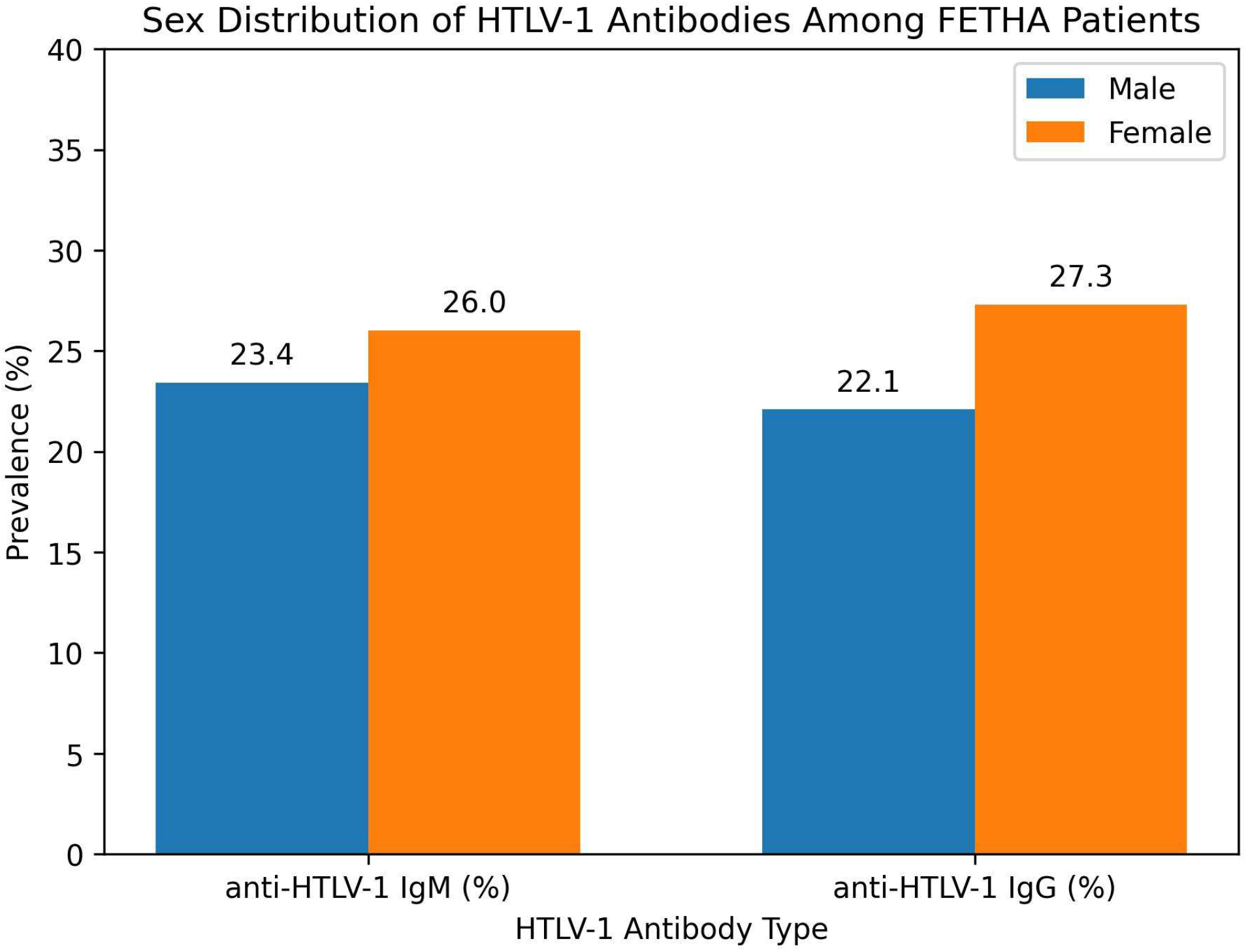
Sex distribution of HTLV-1 IgM and IgG antibodies among patients attending the Federal Teaching Hospital Abakaliki (FETHA), Ebonyi State

Table 2 revealed that the most common was the prevalence of anti-HTLV-1 IgM and IgG antibody with the highest prevalence of 6 (37.5) subjects who gave positive results on both the anti-HTLV-1 IgM and IgG antibodies. The biggest occupational group in the study population was composed of students where 35 (37.6) students were tested to participate in the study. Of them, 9 (25.7) were positive in the anti HTLV-1 IgM antibodies and 8 (22.9) were positive in the anti HTLV-1 IgG antibodies. Moderate prevalence was observed in traders; 5 (23.8%) of the subjects were positive to anti-HTLV-1 IgM and IgG antibodies. Antibodies to HTLV-1 were least common among civil servants where 3 (14.3) and 4 (19.0) had anti-HTLV-1 immunoglobulin (Ig) M and IgG antibodies respectively.

**Table 2:** Occupational Distribution of HTLV-1 Antibodies Among The Study Subjects.

| Occupation | No. of Subjects Tested (%) | Anti-HTLV-1 IgM n (%) | Anti-HTLV-1 IgG n (%) |
| --- | --- | --- | --- |
| Artisan | 16 (17.2) | 6 (37.5) | 6 (37.5) |
| Student | 35 (37.6) | 9 (25.7) | 8 (22.9) |
| Trader | 21 (22.6) | 5 (23.8) | 5 (23.8) |
| Civil Servant | 21 (22.6) | 3 (14.3) | 4 (19.0) |
| **Total (%)** | **93 (100)** | **23 (24.6)** | **23 (24.6)** |

The variance analysis of the serological pattern of infection with the antibodies of HTLV-1 described in the study subjects is presented in Table 3. The values of the different serological patterns differed significantly between one another with the highest mean values observed in those subjects who were positive to both antibodies (IgG+IgM+), and the least mean values were seen with those who were negative of both antibodies (IgG-IgM-). Smaller mean values were observed in the group of the subjects that were positive to IgG only (IgG + IgM -) and to IgM only (IgG - IgM +). In the same way, when comparing the seronegative subjects, there were major variations between the various serological patterns, whose mean values decreasing between IgG-IgM+ to IgG-IgM-. The differences between the values of means were significant as indicated.

**Table 3:**
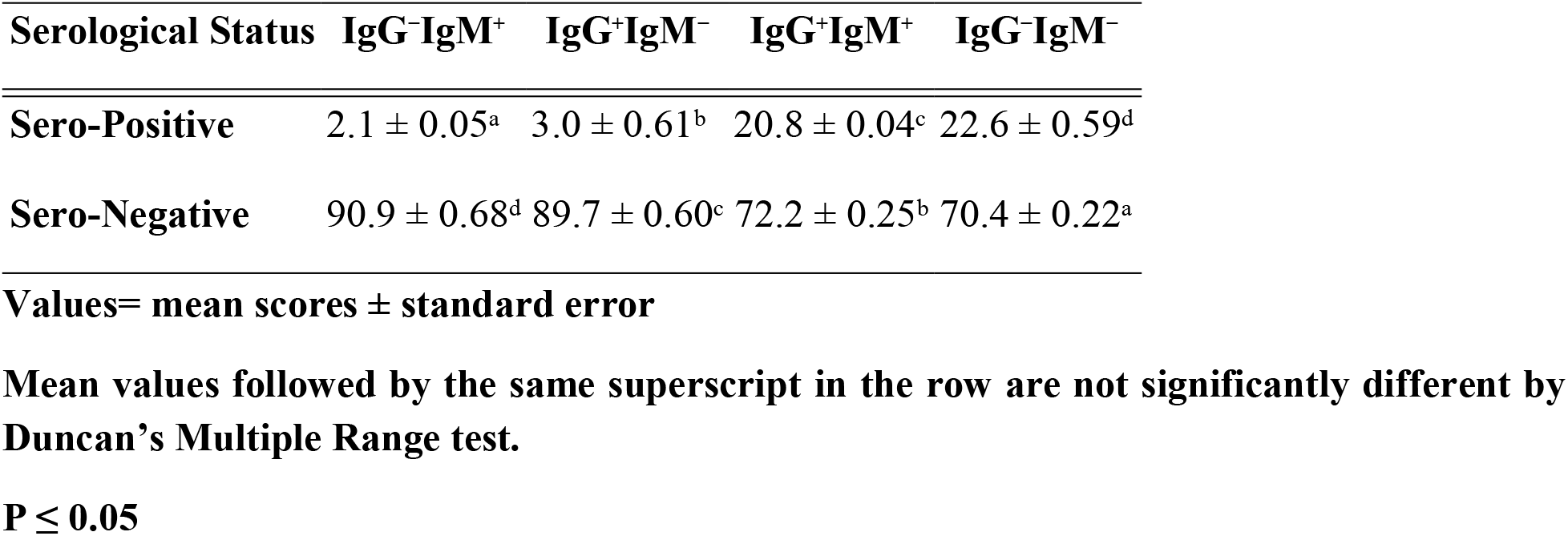
Variance Analysis for Serological Pattern of HTLV-1 Antibodies among Subjects Studies.

## Discussion

This paper showed that 23 out of 93 qualified blood donors (24.6) were seropositive to both HTLV-1 IgM and IgG antibodies by ELISA. This is also a significantly high prevalence as compared to most blood donor studies in Nigeria and sub-Saharan Africa, where the prevalence of HTLV-1 is generally low (usually less than 1 percent to several percent) by geography, donor selection, and assay strategy. A systematic review of the HTLV seroprevalence in blood donors in sub-Saharan Africa stated that there were overall prevalence rates that were low and varied within countries and design but was noted that estimates were heterogeneous and could be attributable to epidemiologic variation and testing methods [12].

In Nigeria, other studies that are based on donors have also indicated lower seroprevalence than those in the current research. As an illustration, the University of Nigeria Teaching Hospital (Enugu) donor study (preferred seroprevalence) reported 0% seroprevalence in their study [13], but another study of donor prevalence in Nigeria has incorporating confirmatory testing (e.g., Western blot or PCR), replacement-donor profile of most tertiary hospitals, and variation in the risk profile of the donors themselves [14]. These points have been continuously emphasized in the literature on transfusion-transmitted infection, in which screening policies and confirmatory testing have significant impact on reported prevalence and consequences of donor deferral.

The identified age-specific trend, where age groups of middle adults showed the greatest positivity, is not unique to studies on HTLV-1, with the prevalence of the disease often positively correlated with age due to accumulated exposure over time (sexual transmission, previous parenteral exposures, and other risk factors) [15,16]. The distribution of sexes in this study indicated that there was a slight prevalence of the sex distribution between female and male donors. This observation is in agreement with the literature which suggests that male-to-female sexual HTLV-1 transmission is generally more effective compared to female-to-male transmission, which helps explain sex differences in seropositivity [17]

More seropositivity was observed among married donors, which can be linked to sexual transmission as a significant method of HTLV-1 transmission [18]. In the context where regular screening of HTLV-1 is not established, the undiagnosed chronic carriers may be able to perpetuate the process of transmission in steady relationships, particularly in situations with low awareness and lack of conventional preventive counseling. Recent literature on the topic of the HTLV screening programs contends that blood donor screening, though valuable, can be inadequate to achieve larger objectives in transmitting reduction to the population with integrated prevention strategies.

Among artisans, occupational distribution demonstrated the greatest seroprevalence when compared to civil servants, traders and students. Occupation may not be causal, but it may serve as a proxy of socioeconomic status, patterns of healthcare access, and risk exposures of behavior. Other Nigerian and African sero-epidemiologic studies have been done with similar interpretations using occupational categories that are associated with differences in mobility, medical injection exposure, informal medical care use and other contextual risk factors [19,20].

The occurrence of HTLV-1 seropositivity among donors supports the debate that continues regarding routine screenings of HTLV in the environment in which screening has not become the norm. Syntheses of evidence in sub-Saharan Africa indicate that prevalence is low on average, but their heterogeneity is high and should be used to inform policy decisions locally[21].

## Conclusion

**T**he paper determined seroprevalence rate and pattern of serology of human T-lymphotropic Virus type-1 (HTLV-1) infection in blood donors in a tertiary health facility in Abakaliki, Ebonyi State, Nigeria. It has been found that the seroprevalence rate of HTLV-1 among eligible blood donors was 24.6 percent as a whole. Both IgM and IgG anti-HTLV-1 antibodies are positive and this indicates that the population of the donors is not only infected recently but also previously. There was a discrepancy in the distribution of antibodies to HTLV-1 regarding demographic characteristics. In addition, seroprevalence was greater among middle-aged donors, those who are married, women and artisans meaning that cumulative exposure, sexual transmission and socio-behavioral factors could form the foundation of the HTLV-1 infection in the context. These findings also show the potential risk of transfusion-transmitted infection of HTLV-1 in the absence of a regular screening of this virus in blood transfusion services. Although it had high seroprevalence through ELISA-based screening, the lack of confirmatory testing is a weakness and may potentially influence the absolute estimate of prevalence. Nonetheless, the reality that the blood donors are positive with the HTLV-1 serums indicates the seriousness of an issue in the community health and the need to come up with improved measures in addressing the blood safety.

## Declarations

### Authors contribution

All the authors contributed to this research work, starting from the beginning of the research to the stage of developing the manuscripts and approved their submission.

